# Effect of Drought On The Germination of Maize Using PEG (Polyethylene Glycol) As A Substitute For Drought Screening

**DOI:** 10.1101/362160

**Authors:** I. A. Dar, Kamaluddin. Z. A. Dar, P.A. Sofi, A.A. Lone

## Abstract

Drought stress is one of the most important environmental factors in reduction of growth, development and production of plants. Germination of each seed is considered as one of the first and most fundamental life stages of a plant so that, the success in growth and yield production is also depending on this stage. To study the effect of PEG stress on germination and early seedling stages on maize, an experiment were laid out at laboratory conditions of Division of Genetics and Plant Breeding SKUAST-K FoA/RRS Wadura. This investigation was performed as factorial experiment under Complete Randomized Design (CRD) with three replications. Polyethylene glycol stress induced in laboratory caused progressive decline in both the parameters across all genotypes with increase in Polyethylene glycol from 0-20% and both the parameters (length of radical and root biomass) had highest value under control.

## INTRODUCTION

Maize (*Zea mays* L.) is an important cereal crop grown all over the world. Also, it is a stable food and commercial crop (Ti-da *et al.*, 2006). On the other hand, drought stress is one of the most important environmental factors in reduction of growth, development and production of plants so, it is urgent to increase maize yields even under the unfavorable conditions. One of the most important abiotic factors limiting plant germination and early seedling stages is water stress brought about by drought and salinity, which are widespread problems around the world. Among the stages of the plant life cycle, seed germination, seedling emergence and establishment are the key processes in the survival and growth of plants (Hadas, 2004). Germination is regulated by duration of wetting and the amount of moisture in the growth medium (Schutz and Milberg 1997; Gill *et al.*, 2002). Water stress acts by decreasing the percentage and rate of germination and seedling growth (Delachiave and De Pinho, 2003). Water stress not only affects seed germination but also increases mean germination time in crop plants (Willenborb *et al.*, 2004).

Solutions of high molecular weight, polyethylene glycol (PEG), are often used to control water potential in seed germination studies (Hardegree and Emmerich, 1990). Polyethylene glycol compounds have been used to simulate osmotic stress effects in petri dish (in vitro) for plants to maintain uniform water potential throughout the experimental period. (Dodd and Donovan, 1999; Sidari et al., 2008). Induced water deficit by polyethylene glycol showed similar values to that observed in the field (Thill *et al.*, 1979), permitting also vigour evaluation. In similar potential ranges, germination pattern may be different between species or even between varieties of the same species (Therios, 1982). Some species, as maize are sensitive to sodium chloride during germination. Polyethylene glycol of high molecular weight range (6000 or above) cannot inter the pores of plant cells (Oertli, 1985) and thus causes cytorrhysis rather than plasmolysis. Polyethylene glycol is also a better choice for imposing low water potential than the often used solute mannitol because mannitol has been shown to be taken up by plant cells and can cause specific toxic effects on growth (Hohl and Schopfer, 1991; Verslues *et al.*, 1998). The principal aim of this study was to investigate the effects of osmotic stress generated by PEG on germination characteristics and early seedling growth of corn.

## MATERIALS AND METHODS

This study was performed at laboratory conditions of Division of Genetics and Plant Breeding SKUAST-K FoA Wadura with thirteen corn cultivar as factorial experiment under completely randomized design (CRD) with three replication. Effect of water stress was induced by different osmotic potential levels [0 (control), 10, 15 and 20 %] of PEG 6000 treatments on germination were studied. Thirteen varieties of maize were evaluated in the present study viz., Shalimar maize composite-4 (C-4), C-6, C-8, C-15, Shalimar maize composite −7 (KDM-72), Kishan Ganga-1 (KG-1), Kishan Ganga-2 (KG-2), Pratap Makka −3 (PM-3), Pratap Makka-4 (PM-4), Pratap Makka-5 (PM-5), Pratap Makka-Chari-6 (PM Chari-6), Aravali Makka-1 (AM-1), Gujrat Makka-6 (GM-6). In each level of stress, sixteen seeds of any cultivar were selected and sterilized in sodium hypochlorite (1%) and then washed in distilled water for two times. The seeds of corn were germinated in Petri dishes on 2 layers of filter paper in germinator maintained at 25°C and 75 % humidity in darkness. Daily the need for, to replace the filter papers and add the PEG soluble was performed. After 7 days, germination percent was measured by International Seed Testing Association (ISTA) standard method. At the end of the seventh day, the length of radicle (cm) and radicle weight (g) of seeds were measured.

## RESULTS

On the basis of analysis of variance results, the effect of stress levels on length of radicle (cm) and radicle weight (g) were significant. Mean comparison results also revealed that the length of radicle and radicle weight under different stress levels were different.

1) **Length of Radicle (cm):** The data on length of radical under different levels of PEG-6000 showed following results (Table 1): **Table 1:**
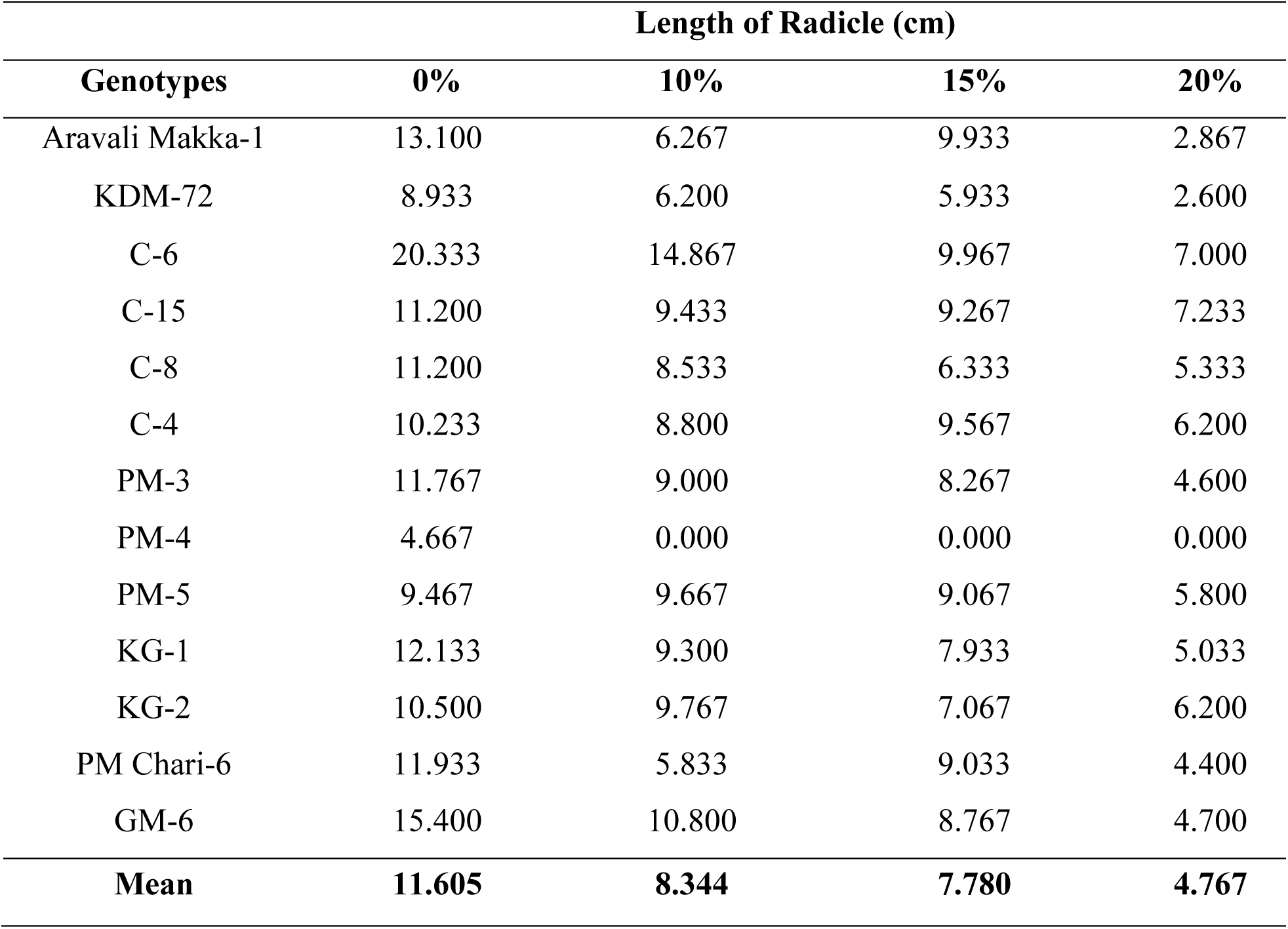
Effect of osmotic potential induced by PEG 6000 on length of radicle (cm) of thirteen maize genotypes

a. **0% Level:-** Under controlled conditions the length of radical was found highest in C-6 (20.33) followed by GM-6 (15.40) and AM-1 (13.10) and was lowest in PM-4 (4.66).
b. **10% Level:-** Under 10% the length of radical was found highest in C-6 (14.86) followed by GM-6 (10.80) and KG-2 (9.76) and was lowest in PM Chari-6 (5.83).
c. **15% Level:-** Under 15% the length of radical was found highest in C-6 (9.96) followed by AM-1 (9.93) and C-4 (9.56) and was lowest in KDM-72 (5.93).
d. **20% Level:-** Under 20% the length of radical was found highest in C-15 (7.23) followed by C-6 (7.00) and C-4 (6.21) and was lowest in KDM-72 (2.60).

2) **Radicle Weight (g):** The data on radicle weight under different levels of PEG-6000 showed following results (Table 2): **Table 2:**
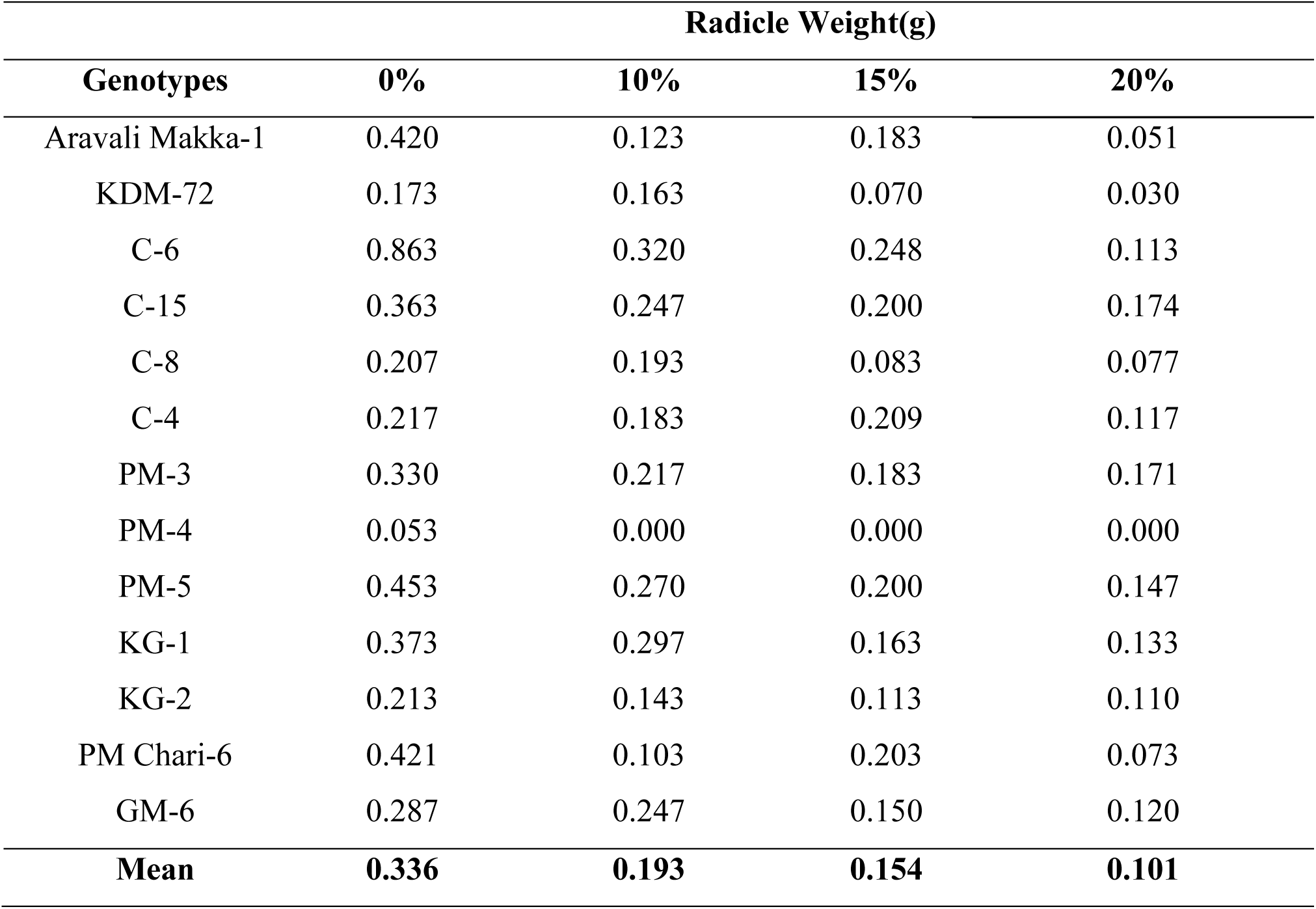
Effect of osmotic potential induced by PEG 6000 on radicle weight (g) of thirteen maize genotypes.

a. **0% Level:-** Under controlled conditions the root biomass was found highest in C-6 (0.863) followed by PM-5 (0.453) and PM Chari-6 (0.421) and was lowest in PM-4 (0.53).
b. **10% Level:-** Under 10% the root biomass was found highest in C-6 (0.320) followed by KG-1 (0.297) and PM-5 (0.270) and was lowest in PM-Chari-6 (0.103).
c. **15% Level:-** Under 15% the root biomass was found highest in C-6 (0.248) followed by C-4 (0.209) and PM Chari-6 (0.203) and was lowest in KDM-72 (0.070).
d. **20% Level:-** Under 20% the root biomass was found highest in C-15 (0.174) followed by PM-3 (0.171) and PM-5 (0.147) and was lowest in KDM-72 (0.030).

## Discussion

Polyethylene glycol is the best solute that we are aware of for imposing a low water stress that is reflective of the type of stress Imposed by a drying soil (Verslues and Bray, 2004; Verslues *et al.*, 1998; Van der Weele *et al.*, 2000). Water stress due to drought is probably the most significant abiotic factor limiting plant and also crop growth and development. Drought stress is physiologically related, because it induces osmotic stress and most of the metabolic responses of the affected plants are similar to some extent. Water deficit affects the germination of seed and the growth of seedlings negatively. In present study, polyethylene glycol (PEG) compounds have been used to stimulate osmotic stress effects in Petri dish (*In vitro*) for plants to maintain uniform water potential throughout the experimental period. There was a progressive decline in both the parameters across all genotypes with increase in PEG levels from 0-20% and both the parameters (Length of radical and radicle weight) had highest value under control (0 %) and had significant differences under water stress (Fig. 1 & 2). The root length provides an important clue to the response of plants to drought stress. Water stress acts by decreasing the percentage and rate of germination and seedling growth as reported by Delachiave and De Pinho (2003) in senna, Farsiani and Ghobadi (2009) and Khayatnezhad et al. (2010) in corn, Gholamin and Khayatnezhad (2010) in wheat, and Mostafavi (2011) in safflower. There were reports in the literature of potential drought resistance traits like extensive viable rooting system that could explore deeper soil layers for water (Mirza, 1956; Bocev, 1963). Corn plants with more roots at seedling stage subsequently developed stronger root system, produced more green matter and had higher values for most characters determining seed yield (Bocev, 1963). This study strongly supports the assertion that germination indices can be utilized to screen maize for drought tolerance at germination and early seedling growth stage. There are many reports that are in agreement with the present findings indicating that drought stress severely reduces seed germination and early seedling growth. But the varieties having genetic potential to maintain the higher growth under stress conditions are drought tolerant. Water stress due to drought is probably the most significant abiotic factor limiting plant and also crop growth and development (Hartman et al., 2005). Growth of plants in arid and semi-arid land is dependent upon plants susceptibility to drought stress and also related to the ability of seeds to achieve optimum germination under these unfavorable conditions. Therefore, it is necessary to identify genotypes that are tolerance to drought at the primary growth stage.

**Fig. 1:**
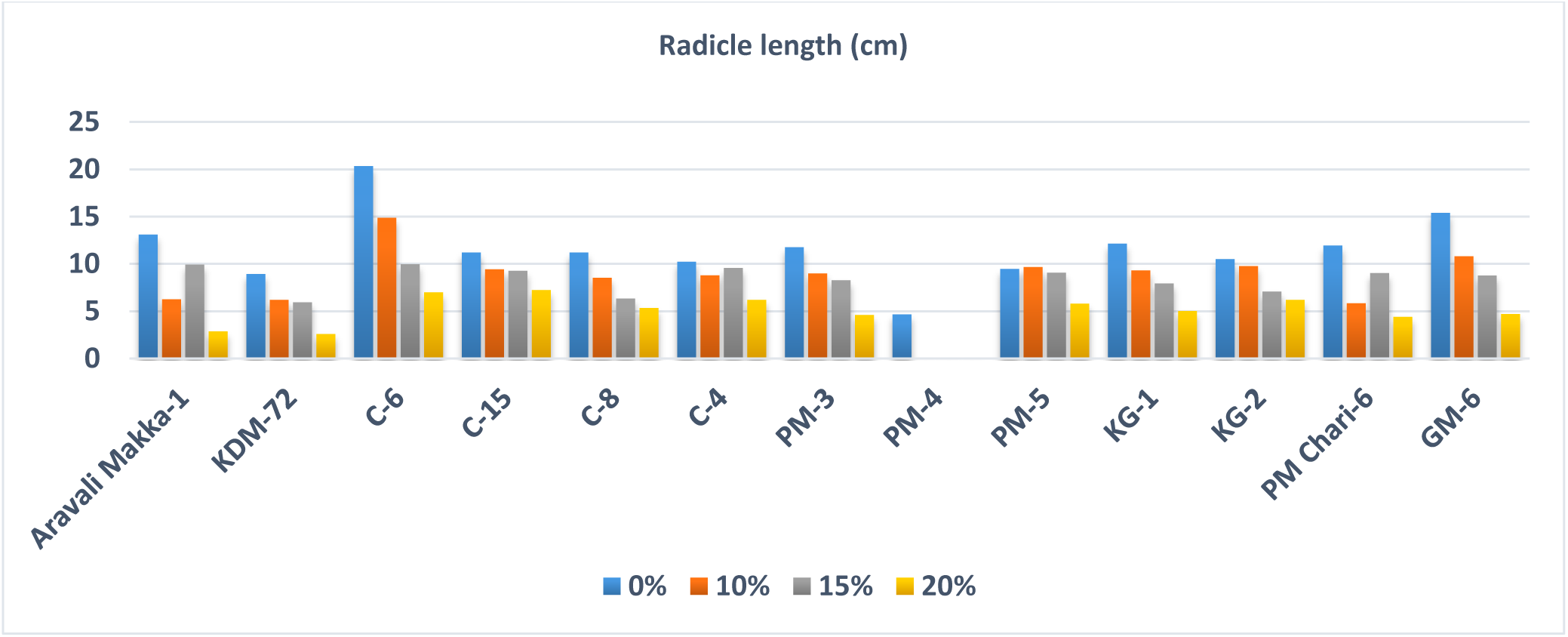
Percentage of length of radicle affected by PEG.

**Fig. 2:**
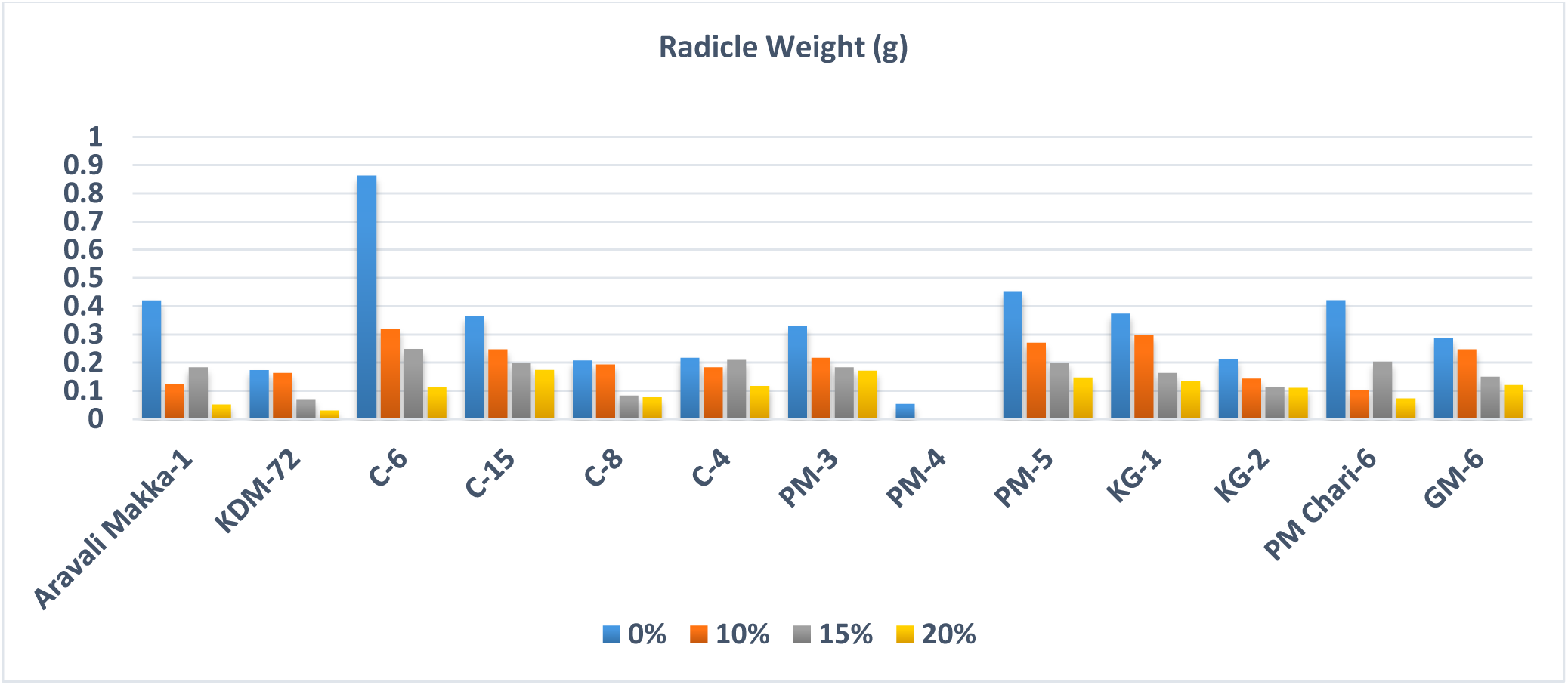
Percentage of radicle weight affected by PEG.

